# Gut microbiota structure differs between honey bees in winter and summer

**DOI:** 10.1101/703512

**Authors:** Lucie Kešnerová, Olivier Emery, Michaël Troilo, Joanito Liberti, Berra Erkosar, Philipp Engel

## Abstract

Adult honey bees harbor a specialized gut microbiota of relatively low complexity. While seasonal differences in community composition have been reported, previous studies have focused on compositional changes rather than differences in absolute bacterial loads. Moreover, little is known about the gut microbiota of winter bees, which live much longer than bees during the foraging season, and which are critical for colony survival. We quantified seven core members of the bee gut microbiota in a single colony over two years and characterized the community composition in 14 colonies during summer and winter. Our data shows that total bacterial loads substantially differ between foragers, nurses, and winter bees. Long-lived winter bees had the highest bacterial loads and the lowest community α-diversity, with a characteristic shift towards high levels of *Bartonella* and *Commensalibacter*, and a reduction of opportunistic colonizers. Using gnotobiotic bee experiments, we show that diet is a major contributor to the observed differences in bacterial loads. Overall, our study reveals that the gut microbiota of winter bees is remarkably different from foragers and nurses. Considering the importance of winter bees for colony survival, future work should focus on the role of the gut microbiota in winter bee health and disease.

## Introduction

The European honey bee, *Apis mellifera,* is an important pollinator species for natural ecosystems and agricultural production [1]. Its health status is threatened by numerous factors including habitat loss, pesticide exposure, and high parasite and pathogen loads [2–4]. Accumulating evidence suggests that the gut microbiota of adult honey bees plays a critical role for bee health [5]. The bee microbiota converts dietary compounds [6, 7] and produces short chain fatty acids [8] in the gut, enhances sucrose responsiveness of the host [8], and stimulates the immune system [9, 10]. Moreover, disruption of the gut microbiota composition by antibiotic treatment, pesticide exposure, or dietary manipulations has been associated with increased pathogen loads resulting in increased host mortality [11–14].

A striking feature of the honey bee gut microbiota is its low taxonomic complexity. In worker bees, the community is dominated by less than ten phylotypes (i.e. clusters of strains sharing ≥97% sequence identity in the 16S rRNA gene), which typically make up >95% of the bacterial cells in the gut [5, 15–18]. These phylotypes have been consistently detected in honey bees, regardless of geographic location, life stage, or season [16, 19, 20], and are acquired horizontally through contact with nest mates and hive components [21]. They include five core phylotypes (*Gilliamella, Snodgrassella, Lactobacillus* Firm-4 and Firm-5, and *Bifidobacterium*), which are typically present in every adult worker bee, and a number of non-core phylotypes, e.g. *Frischella*, *Bartonella, Commensalibacter*, or *Bombella*, which are prevalent across colonies, but not necessarily present in every bee [22]. Additional phylotypes have been detected, including *Lactobacillus kunkeii, Serratia marcescens* and other Enterobacteriaceae, or *Apibacter*, but they typically account for a relatively small proportion of the bee gut microbiota [23].

While many of these phylotypes are consistently present in adult worker bees, their abundance can vary across bees, and may differentially impact the host physiology. Particularly, the type and amount of nutrients (i.e. pollen and nectar) available during the foraging season can have profound effects on the composition of the gut microbiota and may alter its metabolic activity [24]. Likewise, distinct dietary habits or variation in lifespan of worker bees during summer and winter may influence gut microbiota composition. From spring to autumn, young worker bees (nurses) stay inside the hive to take care of larvae, and feed on nutrient-rich pollen, whereas older worker bees become foragers that feed on nectar and honey to fuel their energy-expensive flights [25]. In late autumn, newly emerged adult worker bees become winter bees (also called ‘diutinus’) that have an extended lifespan (∼6 months) and that ensure the colony survival during the cold winter season in the absence of brood [26]. These bees form a tight cluster for thermoregulation inside the hive, feed strictly on food stores (pollen, beebread, and honey) and retain their feces all winter [27], which is likely to impact the ecology of their gut microbiota.

A number of studies have looked at the gut microbiota composition of different worker bee types or throughout seasons, with the overall conclusion that the community composition is relatively stable [19, 28–31]. However, previous studies were mostly based on comparative analyses of relative community member abundance using 16S rRNA amplicon sequencing. Such analysis cannot provide insights about the extent or directionality of changes in taxa abundance, especially if microbial loads vary substantially between samples [32]. In fact, a change in the total abundance of the microbiota could by itself be an important characteristic of different bee types (e.g. foragers, nurses, winter bees), season, or environmental exposure. An example is the experimental exposure of bees to antibiotics which did not result in a strong shift in the relative composition, but in an overall reduction of bacterial load, rendering bees more susceptible to pathogen invasion [11]. In addition to the limitations of current studies using relative abundance data, almost nothing is known about the gut microbiota of winter bees as compared to foragers or nurses. This is surprising, as winter bees are critical for colony health and survival during the cold season of the year, when resources are limited and most colony losses occur [33, 34].

Characterizing the gut microbiota of winter bees and identifying factors that shape its community composition may help to understand the physiological adaptations that honey bees need to survive the cold season in temperate regions. In this study, we used qPCR and 16S rRNA gene amplicon sequencing to assess differences in the gut microbiota of nurses, foragers, and winter bees. We analyzed bacterial loads of major community members in 566 individual worker bees sampled from a single hive over two years. We then expanded our analysis to the entire community and analyzed pooled samples from 14 different hives to test if similar community changes occur in winter bees across hives. Finally, we performed experiments with gnotobiotic bees to test the influence of diet on differences in gut microbiota composition. Our study reveals major differences in total bacterial load and in the abundance of specific gut community members in the gut microbiota of nurses, foragers, and winter bees and identifies dietary pollen as a major contributing factor.

## Materials and Methods

### Sampling of honey bees

Over a period of two years, we sampled *∼*24 adult worker bees of *A. mellifera* each month from a single hive located on the Dorigny campus of the University of Lausanne, Switzerland. These bees were used to determine seasonal changes in the absolute abundance of seven major community members of the honey bee gut microbiota using qPCR. During the foraging season, we sampled foragers returning to the hive entrance with pollen on their legs, while during the cold winter months, we sampled winter bees on top of the frames from inside the hive. Each sampling time point took place at the middle of each month (+/- 3 days) between April 2015 and April 2017. Samples from July 2015 were not included in the analysis due to an error that occurred during DNA extraction.

To identify changes in gut microbiota composition between adult worker bees in summer (i.e. nurses and foragers) and in winter (i.e. winter bees) across colonies, we sampled bees from 14 different hives. Eleven hives were located on the Dorigny campus of the University of Lausanne, and three hives were located in the village of Yens, about 17 km away. Foragers and nurses were sampled in July 2017 and August 2018, and winter bees were sampled in January 2018 and January 2019. Nurses and winter bees were collected inside the hive in the center of the colony. Foragers were collected outside the hive at the same time point as nurses. The bees sampled in July 2017 and January 2018 were used for gut content visualization and correlation of gut weight and 16S rRNA gene copy numbers. The guts of the bees sampled in August 2018 and January 2019 were pooled (20 guts per bee type per hive) and used to monitor the abundance of individual community members by qPCR and to determine the overall community structure by 16S rRNA community analysis.

Bees were anesthetized with CO_2_ for 10 s and the gut including crop, midgut, hindgut, and malpighian tubules carefully removed using sterile forceps. For the monthly sampling of the single bee hive, each gut sample was placed in a drop of PBS, scored for the scab phenotype [35], and placed in a bead beating tube containing *∼*150 mg of glass beads (0.75-1 mm in diameter, Carl Roth) and 750 µL of CTAB lysis buffer (0.2 M Tris-HCl, pH 8; 1.4 M NaCl; 0.02 M EDTA, pH 8; 2% CTAB, w/v, dissolved at 56°C overnight; 0.25% β-mercaptoethanol, v/v). For the sampling of the 14 colonies in August 2018 and January 2019, the dissected guts were pooled into a single Falcon tube (14 hives x 3 bee types = 42 pooled samples). Tubes were flash frozen in liquid nitrogen and stored at −80°C until DNA extraction.

### Experimental colonization of honey bees

Microbiota-depleted bees were generated and colonized as described in Kešnerová et al. [6]. The treatment group of bees fed on pollen and sugar water was the same as in the previous study. The other treatment group (bees fed on sugar water only) was carried out in parallel with the same batch of bees. Bees were fed *ad libitum* with sterilized bee pollen (P) and sterilized sugar water (SW, 50% sucrose, w/v) (SW+P treatment), or with only sterilized sugar water (SW treatment). Bees were sampled 10 days after colonization and the guts were dissected as described before.

### DNA extraction from honey bee gut tissue

A previously established CTAB/phenol-based extraction protocol [6] was used to extract DNA from individual guts. At the end of the protocol, the precipitated dried pellet was resuspended in 200 μl and split into two samples of 100 μl each. One sample was processed with the Nucleospin PCR Clean-up kit (Macherey-Nagel, Germany) according to the manufacturer’s instructions and the resulting DNA was used for qPCR. For the pooled gut samples, 2 ml of glass beads and 15 ml of CTAB lysis buffer were added to each Falcon tube. Samples were then homogenized in a Fast-Prep24™5G homogenizer at 6 m/s for 40 s, briefly centrifuged, and an aliquot of 750 μl corresponding to the sample volume of one bee gut was transferred to a new 2 ml bead beating tube with glass beads and homogenized again. All further steps of the DNA extraction were performed as previously described [6].

### Quantitative PCR (qPCR) to determine absolute abundance of community members

Bacterial absolute abundances were determined using qPCR assays targeting the 16S rRNA gene of either specific community members or universally all bacteria, and normalized to the number of host actin gene copies, as described in Kešnerová et al. [6].

Standard curves were performed on serial dilutions containing known quantities of plasmid DNA encoding the target sequence as follows: The plasmid copy number was calculated based on the molecular weight of the plasmid and the DNA concentration of the purified plasmid. Dilutions containing 10^1^–10^7^ plasmid copies per μl were used to generate the standard curves. For *Frischella*, *Snodgrassella*, *Bartonella*, *Lactobacillus* Firm4, and *Lactobacillus* Firm5, the slope and intercept of the standard curve was calculated based on the Cq values (quantification cycle [36]) obtained from the dilutions containing 10^2^ - 10^7^ plasmid copies. For these targets, the Cq value corresponding to 10^2^ copies was set as the limit of detection (LOD) of the primer set, because dilutions containing 10^1^ copies resulted in Cq values which could not be discriminated from the water control, or the signal was undetected. For all other targets (*Gilliamella*, *Bifidobacterium*, and actin), the slope and intercept of the standard curve was calculated based on the Cq values obtained from all seven dilutions, and the Cq value corresponding to 10^1^ copies was used as the LOD. Bacterial targets resulting in Cq values higher than the LOD of a given primer pair were considered to be too low to be quantified in the respective sample (i.e. <10^2^ or <10^1^ copies per μl). The *Efficiency of primers (E)* was estimated from the slope according to the equation: *E* = 10^(−1/slope)^ [37]. Primer characteristics and their performance are summarized in **Table S1**.

One individual gut sample had to be excluded from the analysis, because it gave no amplification for any of the bacterial targets and very low amplification for actin. For all other samples we determined the number of bacterial genome equivalents per gut as a proxy for bacterial abundance as follows: We first calculated the ‘raw’ copy number (*n_raw_*) of each target in 1 μl of DNA (the volume used in each qPCR reaction) based on the Cq value and the standard curve using the formula *n_raw_* = *E* ^(intercept - Cq)^ [38]. Then, we normalized the ‘raw’ copy number by dividing by the number of actin gene copies present in the sample (*n_actin_*), which was determined using the same formula. This normalized value of 16S rRNA gene copies was then multiplied by the median number of actin gene copies of the samples of a given dataset and the total volume of extracted DNA (i.e. 200 μl) to obtain normalized copy numbers per gut (*n_abs_*): *n_abs_= (n_raw_/n_actin_) x median(n_actin_) x 200.* Normalization with the actin gene was done to reduce the effect of gut size variation and the DNA extraction efficiency. To report the number of genome equivalents (*n_GE_*) rather than the normalized 16S rRNA gene copy number, we divided *n_abs_* by the number of 16S rRNA loci present in the genome of the target bacterium (as listed in **Table S1**). For the qPCR results obtained with the universal bacterial primers, we reported the absolute 16S rRNA gene copies (*n_abs_*) rather than genome equivalents (*n_GE_*), as the number of 16S rRNA gene loci differs between bacteria.

### 16S rRNA gene amplicon sequencing

The V4 region of the 16S rRNA gene was amplified as described in the Illumina 16S metagenomic sequencing preparation guide (https://support.illumina.com/documents/documentation/chemistry_documentation/16s/16s-metagenomic-library-prep-guide-15044223-b.pdf) using primers 515F-Nex (TCGTCGGCAGCGTCAGATGTGTATAAGAGACAGGTGCCAGCMGCCGCGGTAA) and 806R-Nex (GTCTCGTGGGCTCGGAGATGTGTATAAGAGACAGGGACTACHVGGGTWTCTAAT), which contain the adapter sequences for Nextera XT indexes and the primers for the V4 region of the 16S rRNA gene [39]. PCR amplifications were performed in a total volume of 25 μl, using 12.5 μl of Invitrogen Platinum SuperFi DNA Polymerase Master Mix, 5 μl MilliQ water, 2.5 μl of each primer (5 μM), and 2.5 μl of template DNA. PCR conditions were set to 98°C for 30 s followed by 25 cycles of 98°C for 10 s, 55°C for 20 s and 72°C 20 s, and by a final extension step at 72°C for 5 min. Amplifications were verified by 2% agarose gel electrophoresis. The PCR products were next purified using Clean NGS purification beads (CleanNA) in a 1:0.8 ratio of PCR product to beads, and eluted in 27.5 μl of 10 mM Tris pH 8.5. Next, we performed a second PCR step to add unique dual-index combinations to each sample using the Nextera XT index kit (Illumina). Second-step PCR amplifications were performed in a total volume of 25 μl using 2.5 μl of the PCR products, 12.5 μl of Invitrogen Platinum SuperFi DNA Polymerase Master Mix, 5 μl MilliQ water, and 2.5 μl each of Nextera XT index primers 1 and 2. Thermal cycle conditions were an initial denaturation step at 95°C for 3 min followed by eight cycles of 30 s at 95°C, 30 s at 55°C, and 30 s at 72°C, and a final extension step at 72°C for 5 min. The final libraries were purified using Clean NGS purification beads in a 1:1.12 ratio of PCR product to beads, and eluted in 27.5 μl of 10 mM Tris pH 8.5. The amplicon concentrations, including the negative PCR control, were then quantified by PicoGreen and pooled in equimolar concentrations (with the exception of the negative control). We verified that the final pool was of the right size using a Fragment Analyzer (Advanced Analytical) and performed sequencing on an Illumina MiSeq sequencer, producing 2 x 250 bp reads, at the Genomic Technology Facility of the University of Lausanne.

### Processing of 16S rRNA gene amplicon sequencing data

Divisive Amplicon Denoising Algorithm 2 (DADA2) pipeline (“dada2” package version 1.12.1 in R) was used to process the sequencing data (see script ‘2_Dada2_Pipeline.R’ on Zenodo) [40]. All functions were ran using the recommended parameters (https://benjjneb.github.io/dada2/tutorial.html) except for “expected errors” during the filtering step which was set to (maxEE=1,1) in “filterAndTrim” function. The SILVA database was used for taxonomy assignments. Downstream analyses were performed in R version 3.6.0. Reads belonging to mitochondria, chloroplast, and eukaryotes were excluded from further analyses (“phyloseq” package version 1.28.0 [41], “subset_taxa” function). Only reads that are present in at least two samples with a total number of 10 reads were retained for downstream analyses (“genefilter” package version 1.66.0 [42], “filterfun_sample” function, see script ‘2_Dada2_Pipeline.R’ on Zenodo). To complement the taxonomic classification based on the SILVA database, sequence variants were further assigned to major phylotypes of the bee gut microbiota as defined in previous studies based on a BLASTn search against the Nucleotide (nt) database of NCBI (https://www.ncbi.nlm.nih.gov/nucleotide/). To analyze absolute bacterial abundances, we multiplied the proportions of each taxon by the total 16S rRNA gene copy number present in each sample (as measured by qPCR using the universal bacterial primers and normalized by actin copy gene number), and divided this number by the number of 16S rRNA loci for each taxon. The mean 16S rRNA operon copy number for each taxon was obtained from a previous study [11] and completed from rrNDB (https://rrndb.umms.med.umich.edu/).

### Diversity analysis and statistics

Diversity analyses were performed using “Vegan” package [43]. For both datasets, the qPCR data from the monthly sampling and the 16S rRNA gene amplicon data from the pooled samples, we measured *α*-diversity using effective number of species [44] that is calculated by taking the exponent of Shannon’s diversity index (“diversity” function). For the 16S rRNA gene amplicon sequencing data, permutational multivariate analysis of variance (ADONIS, “adonis” function) based on Bray–Curtis dissimilarities (“vegdist” function) [45] was used to test the effect of bee type on community structure, and “metaMDS” function was used for plotting beta-diversity (see script ‘3_Plots_Stats_Figures2_S4.R’ on Zenodo). To test the dispersion of communities we used the function “betadisper” [46, 47] and compared the distances of individual samples to group centroids in multidimensional space using “permutest”. For the qPCR data from the monthly sampling, we performed a principal component analysis (PCA) with the *prcomp* function of the R package “stats” to determine the similarity of the bacterial communities between foragers and winter bees using absolute abundance measures of the seven gut microbiota phylotypes.

All statistical analyses were performed using R (version 3.6.0). We tested the effect of bee type on bacterial loads, diversity indices, and wet gut weight using Student’s t-test (in case of two group comparisons) or general linear models (in case of three group comparisons). Since the residuals obtained for certain models showed heteroscedasticity, we used a permutation approach (referred to as Permutation TTEST or ANOVA respectively) to test the significance of the effects as described before [48]. Briefly, we randomized the values of the response variable 10,000 times and computed the F-values/t-values for the tested effect for each randomized dataset. The p-values corresponding to the effects were calculated as the proportion of 10,000 F-values that were equal or higher than the observed one. Pairwise comparisons between different factors were performed by Tukey’s HSD using “multcomp” package [49] using *glht* function on the model. P-values were adjusted using the Bonferroni method. Detailed results of statistics are reported in Supplementary tables S2-S5.

## Results

### Bacterial loads of core microbiota members differ between foraging and winter season in a honey bee colony monitored over two years

To characterize the gut microbiota of adult worker bees across seasons, we tracked the total abundance of five core (*Gilliamella*, *Snodgrassella, Bifidobacterium*, *Lactobacillus* Firm4, and *Lactobacillus* Firm5) and two non-core members (*Frischella,* and *Bartonella*; **Table S1**) in adult worker bees from a single hive over two years. Our analyses included 566 individual bee samples.

The core members *Gilliamella*, *Snodgrassella*, *Lactobacillus* Firm-5 and *Bifidobacterium* were present in all analyzed bees, and the core member *Lactobacillus* Firm-4 was detectable in 98.4% of all bees (**Supplementary Fig. S1A**). Notably, the two designated non-core members *Bartonella and Frischella* were also present at relatively high prevalence with only 5.3% and 26.9% of the samples giving signals below the detection limit, respectively (**Supplementary Figs. S1B** & **C**). Consistent with our previous results, *Frischella* prevalence strongly correlated with the presence of the scab phenotype (**Supplementary Fig. S2**), a local melanization response that is induced by *Frischella* upon colonization [35].

The absolute abundance of the monitored phylotypes varied little among the bees sampled at the same time point, with the exception of the non-core members *Frischella* and *Bartonella* (**Supplementary Fig. S3**). However, there were clear differences in bacterial abundances between months (Permutation ANOVA P=1e-4) for all monitored community members. In particular, we observed remarkable differences in the bacterial loads between bees sampled during the foraging and the winter season in both years. This became evident from the abundance of individual phylotypes and from the total bacterial load inferred from the summed abundances of all seven phylotypes (**Supplementary Fig. S3,** Fig. 1A). Specifically, we found a 10- to 100-fold increase in the levels of the core members *Lactobacillus* Firm-4, *Lactobacillus* Firm-5, and *Bifidobacterium*, as well as the non-core member *Bartonella* when comparing across all winter bees relative to foragers (Fig. 1C, Permutation T-Test P=1e-4). We also observed a small increase of *Snodgrassella* levels in winter bees (Fig. 1C, Permutation T-Test P=6e-4), but no difference in the levels of *Gilliamella* (Fig. 1C, Permutation T-Test P=0.7). *Frischella* was the only member of the community that displayed the opposite trend, i.e. lower abundance in winter bees (Fig. 1C, Permutation T-Test P=1e-4). The overall bacterial load was about 10x larger in winter bees than in foragers based on both the summed abundances of all seven phylotypes (Fig. 1C, Permutation T-Test P=1e-4) as well as the number of total 16S rRNA gene copies, which was determined with universal 16S rRNA gene qPCR primers for a subset of the samples (Fig. 1D, Permutation T-Test, P=1e-4).

**Fig. 1.**
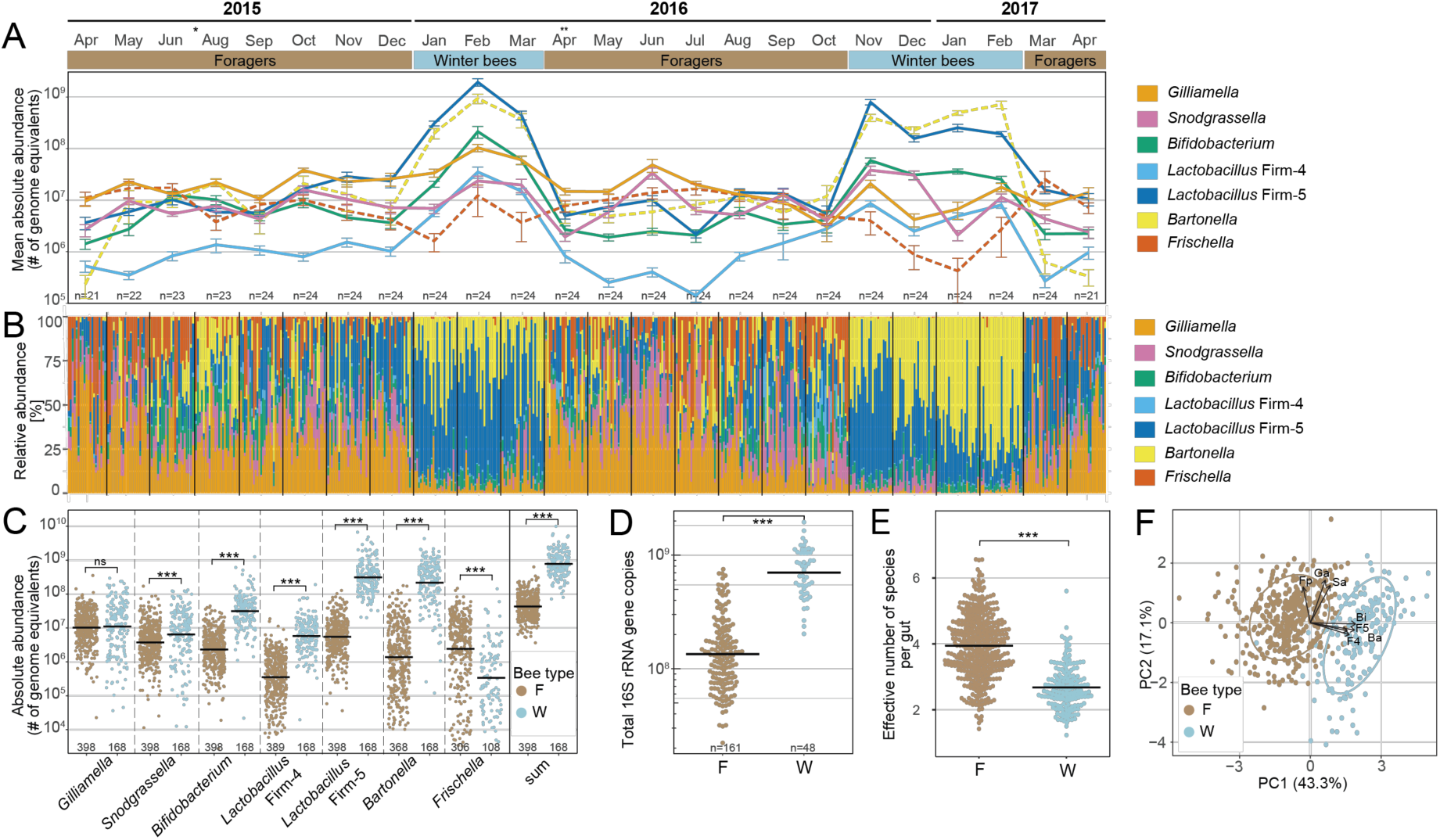
Gut bacterial communities differ between foraging and winter season in a single colony monitored over two years. **(A)** Monthly changes in the absolute abundance assessed by qPCR, as determined by number of genome equivalents per sample, of seven phylotypes monitored over a period of two years, depicted as mean values (±SE) of the analyzed bees. The number of bees per month is indicated at the bottom of the plot. * indicates missing data for July 2015 due to DNA extraction failure. ** indicates that the queen of the colony was replaced in the corresponding month. **(B)** Relative community composition of the gut microbiota of bees sampled in each month as based on the seven monitored phylotypes. **(C)** Absolute bacterial abundance of each phylotype per gut in foragers (F) and winter bees (W), as determined by the number of genome equivalents. The sum of the abundances of the seven monitored phylotypes is also plotted. Mean values are shown as black horizontal lines. Only bees with detectable levels were plotted (the number of bees is given at the bottom of the plot; for prevalence see **Supplementary Fig. S1**). **(D)** Copy number of the 16S rRNA gene in gut samples of a subset of the analyzed months (Apr 2015, Aug 2015, Oct 2015, Jan 2016, Apr 2016, Jul 2016, Oct 2016, Jan 2017, Apr 2017). **(E)** Effective number of species calculated from cell numbers of different bacterial phylotypes in foragers (F) and winter bees (W). **(F)** Projection of the abundances of monitored phylotypes into first and second principal components in all analyzed bees, together with correlation vectors representing variables driving the separation on both axes. Permutation T-Test was used for pairwise comparisons. ns, non-significant; ***, P < 0.001.

Considering that the monitored phylotypes typically comprise the majority of the bacteria present in the honey bee gut, we neglected the possible presence of additional, non-targeted members and analyzed the relative composition of the community based on our data. In both years, the communities of winter bees were largely dominated by the phylotypes *Lactobacillus* Firm-5 and *Bartonella.* In contrast, forager bees seem to have more even community compositions (Fig. 1B). The dominance of *Lactobacillus* Firm-5 and *Bartonella* in winter bees was reflected by a reduction in *α*-diversity in winter bees compared to foragers, as determined by the effective number of species (Fig. 1E, Permutation T-Test, P=1e-4). Moreover, PCA revealed a clear separation between foragers and winter bees (Fig. 1F, MANOVA Wilks =0.6 F_(7, 392)_=39.4, P<2.2e-4) along the principal component 1 (PC1). This separation was mainly driven by *Lactobacillus* Firm-4, *Lactobacillus* Firm-5, *Bartonella* and *Bifidobacterium*, the four phylotypes with the largest differences in abundance between the two types of bees (Fig. 1C).

Taken together, these results suggest that the gut microbiota of winter bees and foragers markedly differs from each other in the monitored hive, both in terms of the total bacterial abundance and in the levels of individual microbiota members.

### Consistent difference in bacterial loads and community composition between foragers, nurses, and winter bees across colonies

The observed differences in bacterial loads between foragers and winter bees in the monitored hive prompted us to check for similar patterns across 14 different hives in a subsequent year. In addition to foragers and winter bees, we also analyzed nurses, to help understand whether microbiota differences between foragers and winter bees are linked to seasonal changes or to behavioral or dietary differences. Moreover, we combined our qPCR approach with 16S rRNA gene amplicon sequencing to expand our analysis to the complete community of the honey bee gut microbiota.

Performing universal 16S rRNA qPCR, we found that total bacterial loads differed between the three bee types across the sampled hives. Both winter bees and nurses had higher bacterial loads than foragers (Fig. 2A, Permutation ANOVA P=1e-4, followed by Tukey HSD test, P=5.1e-9 and 1.66e-6 respectively) confirming our previous results from the single hive. Winter bees also showed a trend towards higher bacterial loads than nurses, but this difference was not statistically significant (Fig. 2A, Permutation ANOVA followed by Tukey HSD test, P=0.224). 16S rRNA amplicon sequencing yielded 70 amplicon sequence variants across the 42 samples, with a minimum of 26,993 reads per sample after quality and abundance filtering (See methods and Supplementary Table S6 for details). These sequence variants were further clustered by assigning them to the major phylotypes of the bee gut microbiota as defined in previous studies, resulting in 28 operational taxonomic units (OTUs). To account for the differences in total bacterial load, we calculated absolute abundance of each OTU based on its proportion in the community, the number of rRNA loci in the genome, and the total bacterial load per sample.

**Figure 2.**
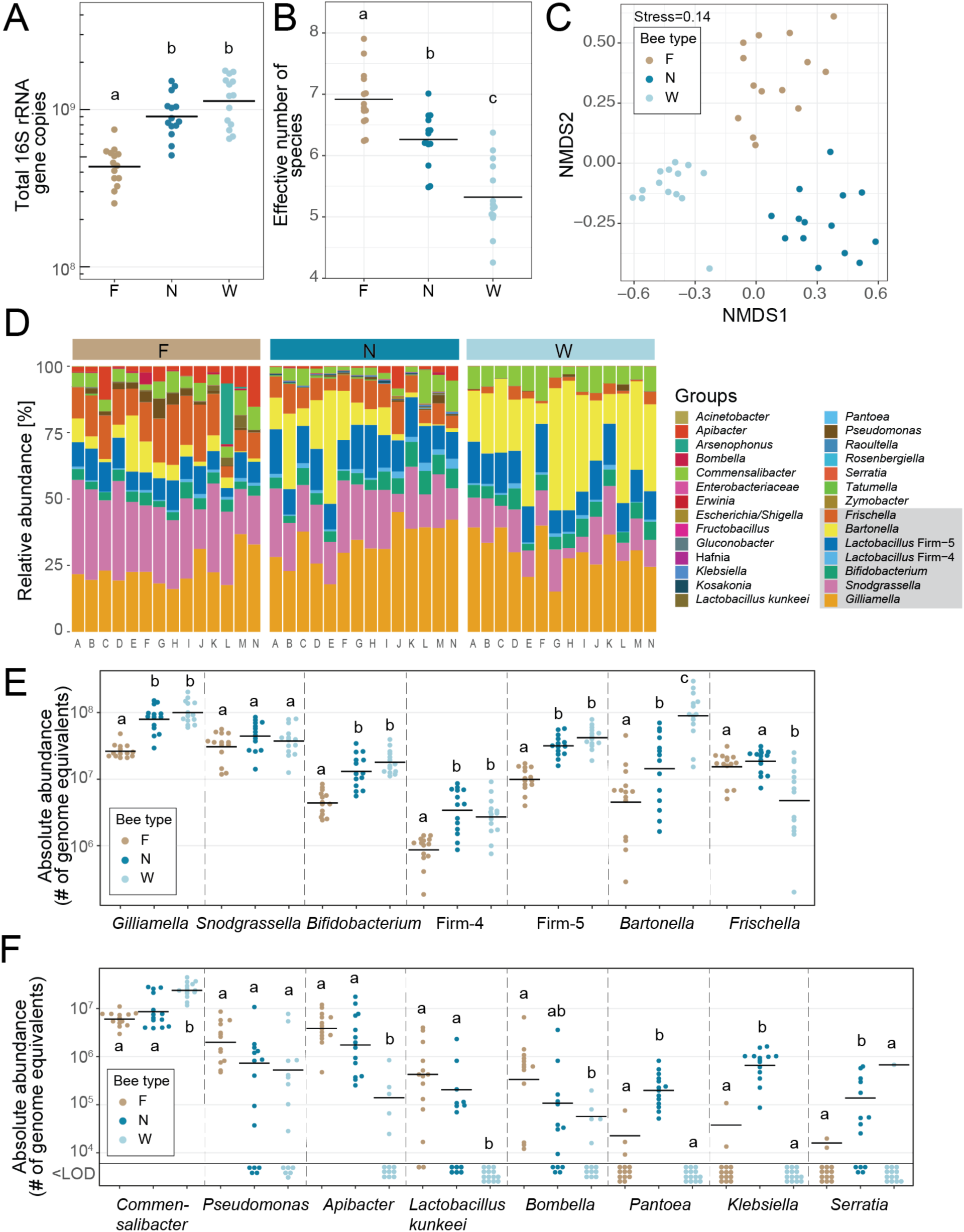
Bacterial load and community composition differ between foragers, nurses, and winter bees across 14 colonies. (**A**) Average 16S rRNA gene copy number per gut in foragers, nurses, and winter bees as determined from the pooled gut samples from the 14 different colonies. **(B)** Differences in *α*-diversity, i.e. effective number of species, in the gut microbiota of foragers, nurses, and winter bees based on 16S rRNA gene amplicon sequencing. **(C)** NMDS based on Bray-Curtis dissimilarities on the gut communities of foragers, nurses, and winter bees based on 16S rRNA gene amplicon sequencing. **(D)** Relative community composition of the gut microbiota based on 16S rRNA gene amplicon sequencing. The seven phylotypes monitored by qPCR (see Fig. 1 and **Supplementary Fig. S4**) which make up the vast majority of the community are highlighted by a grey box in the legend. Capital letters below the stacked bars indicate the hive of origin. **(E)** Absolute abundance of each of the seven major phylotype in foragers (F), nurses (N), and winter bees (W) across hives, as determined based on the number of genome equivalents per gut calculated by multiplying the relative abundance of each phylotype by the total 16S rRNA gene copy number. **(F)** Absolute abundance of a subset of the minor community members in foragers (F), nurses (N), and winter bees (W) across hives, as determined based on the number of genome equivalents per gut calculated by multiplying the relative abundance of each phylotype by the total 16S rRNA gene copy number. <LOD, below limit of detection of the 16S rRNA amplicon sequencing, i.e. no reads were obtained for that particular taxa in the respective sample. Absolute abundance of the remaining phylotypes are depicted in **Supplementary Fig. S4B**. In panels A, B, E and F, levels (bee types) not connected by the same letter are significantly different as based on ANOVA followed by Tukey’s HSD test (see **Supplementary Table S3**).

Diversity analyses of the amplicon sequencing data revealed marked differences in community composition between the three bee types. We found a significant reduction in *α*-diversity in winter bees compared to foragers and nurses, as determined by effective number of species (ANOVA F(2,39) =35.9, p=1.60e-9, Tukey HSD test P<0.005 for all comparisons, Fig. 2B). This indicates that gut communities in these bees are less rich and less even. Moreover, nonmetric multidimensional scaling (NMDS) based on Bray-Curtis dissimilarities revealed a significant separation of samples according to bee type indicating that the communities of nurses, foragers, and winter bees are different from each other (Fig. 2C). Consistently, ADONIS on Bray-Curtis dissimilarities showed a statistically significant difference according to bee type (P=0.001). Differences in community structure were also evident from the relative proportion of different taxa across the samples, with a clear reduction of the relative abundance of *Snodgrassella* and *Frischella* and an increase of *Bartonella* and *Commensalibacter* in winter bees relative to foragers and nurses (Figure 2D). However, we did not detect any difference in community dispersal between nurses, foragers, and winter bees. Distances to group centroids based on Bray-Curtis dissimilarities were not different between bee types (Permutest F_(2,39)_=0.41, P=0.68 (**Supplementary Fig. 4A**). Therefore, while the gut communities of the three bee types differed from each other, they seemed to be similarly variable among each other.

We next assessed differences in the absolute abundance of individual community members to reveal the directionality of community changes. We first looked at the seven phylotypes that were monitored by qPCR over two years (Fig. 2E). Consistent with our previous results (Fig. 1C), *Bifidobacterium*, *Lactobacillus* Firm4, *Lactobacillus* Firm5, and *Bartonella* had increased levels (Permutation ANOVA on three groups P=1e-4, followed by Tukey, P<2e-4), while *Frischella* had decreased levels in winter bees compared to foragers (Fig. 2E, Permutation ANOVA p=2e-04 followed by Tukey HSD test P=2.79e-3). The only two phylotypes showing abundance patterns inconsistent with the results from the two year sampling were *Snodgrassella* and *Gilliamella*. *Snodgrassella* did not experience any differences in absolute abundance (Fig. 2E, Permutation ANOVA P=1.87e-1), illustrating that a proportional change in the community, as found when looking at the relative community composition (Fig. 2D), does not necessarily imply a change in abundance. When comparing nurses and winter bees, only *Bartonella* and *Frischella* showed differences in their absolute abundance. While *Bartonella* had markedly increased levels, *Frischella* abundance went down in winter bees as compared to nurses (Figure 2E). We confirmed these changes by carrying out qPCR on the same samples with the phylotype-specific primers used for the monthly sampling as presented above (**Supplementary Fig. S5**). Except for *Lactobacillus* Firm5, which showed a significant difference between nurses and winter bees in the qPCR but not in the amplicon sequencing data, the results of the two approaches were surprisingly congruent corroborating our conclusion that the microbiota of nurses, foragers, and winter bees markedly differs in the composition of these seven major community members.

We also looked at abundance changes of other community members than those assessed by qPCR the two-year sampling from a single hive (Fig. 2F and **Supplementary Fig. S4B)**. As expected, other bacteria made up a relatively small fraction of the overall community (4-25%) with *Commensalibacter* being the most prevalent (100% of the pooled gut samples) and abundant one (2-14% of the community). *Commensalibacter* was also the only additional community member that showed a significant increase in winter bees compared to foragers and nurses (Permutation ANOVA P=1e-04, followed by Tukey P=7.68e-08). In contrast, all other additional community members were only detected in a subset of the samples, and at relatively low abundance, suggesting that they represent opportunistic or transient colonizers. Moreover, many of these community members showed a trend towards lower prevalence and/or abundance in winter bees than in foragers and nurses (Fig. 2F). For example, while *Apibacter* was detected in all forager and nurse samples, it was only detected in five out of the 14 sampled hives in winter bees and at lower levels than in foragers and nurses (Permutation ANOVA P=1e-04, followed by Tukey HSD test P<2e-8). Likewise, while *Lacotbacillus kunkeii* was detected in nurses and foragers from some hives, it was not detected in any hive during winter. These differences are likely responsible for the reduction in *α*-diversity in winter bees as compared to foragers and nurses. Interestingly, several *Enterobacteriaceae* (*Klebsiella*, *Pantoea*, *Serratia*, or *Tatumella*) were prevalent among nurse samples but absent from nearly all foragers and winter bee samples (Fig. 2F and **Supplementary Fig. S4B**), suggesting a specific association of these bacteria with nurse bees.

Taken together, these results show that winter bees and nurses across hives have increased bacterial loads compared to foragers, and that winter bees have particularly high levels of *Bartonella* and *Commensalibacter*, but low levels of opportunistic colonizers.

### Pollen diet increases gut community size in gnotobiotic bees

One of the drivers of the observed differences in bacterial load and community composition between winter bees, foragers, and nurses could be diet. Dietary differences between the three types of bees were evident from visual inspection of the dissected guts (Fig. 3A-C). The rectums of winter bees and nurses appeared yellow indicating the presence of pollen, while those of foragers were translucent. Moreover, the wet weight of the guts was significantly different between the three types of bees (ANOVA F_(2,68)_ = 24.13, P=1.21e-8), with foragers having on average two times lighter guts than nurses (Tukey HSD test P=1.14e-6) and winter bees (Tukey HSD test P=6.81e-8) (Fig. 3D). When plotting normalized 16S rRNA gene copy numbers as a function of gut weight, we found that gut weight positively correlated with total microbiota abundances across the three bee types (Fig. 3E).

**Figure 3.**
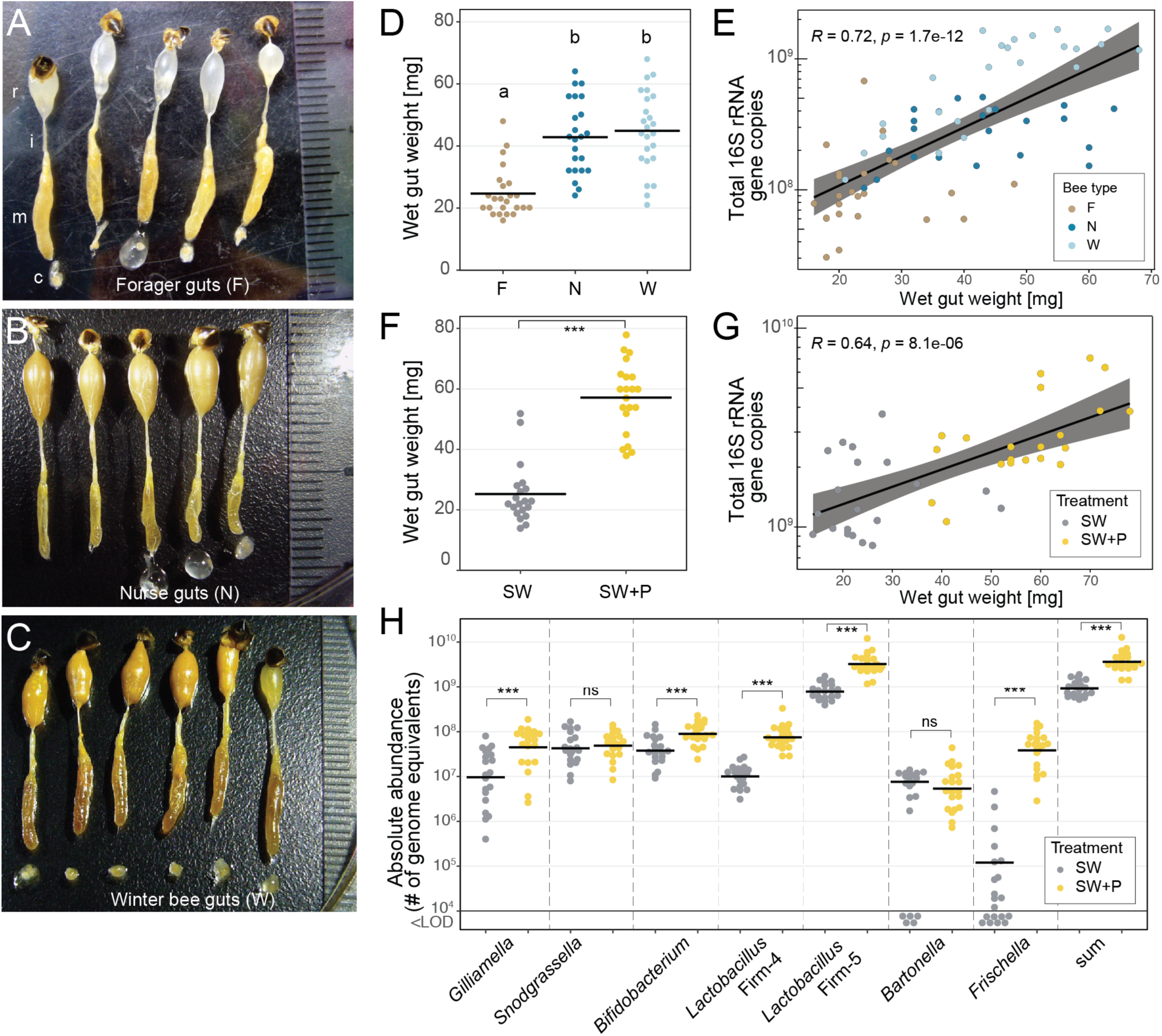
Diet is a major factor shaping the gut microbiota of honey bees. **(A-C)** Dissected guts of foragers **(A)**, nurses **(B)**, and winter bees **(C)** consisting of the crop (c) (missing in some samples), the midgut (m), the ileum (i), and the rectum (r) with attached stinger and last abdominal tergite. **(D)** Wet gut weight of individual foragers (F), nurses (N), and winter bees (W). Different letters indicate groups that are significantly different from each other based on ANOVA followed by Tukey’s HSD test (See Supplementary Table S4 for details). **(E)** Spearman correlation between gut weight and bacterial loads (assessed with universal bacterial 16S rRNA primers) across all bee types. The grey area indicates the 95% confidence interval. **(F)** Wet gut weight of experimentally colonized bees that were fed either sugar water (SW) or sugar water and pollen (SW+P). **(G)** Spearman correlation between gut weight and bacterial loads (assessed with universal bacterial 16S rRNA gene primers) across the two treatment groups of colonized bees. **(H)** Total abundances of the seven monitored phylotypes in the two treatment groups of colonized bees, as determined by genome equivalents per gut using phylotype-specific qPCR primers. The sum of the abundances of the seven monitored phylotypes is also depicted. Bees with bacterial loads below the limit of detection (<LOD) of the qPCR method are shown below the cut of the axis at 10^4^. Two-group comparisons were done by Permutation T-Test. ns, non-significant; ***, P < 0.001.

In order to demonstrate that pollen diet is directly associated with increased bacterial loads in honey bees, we experimentally colonized newly emerged bees with a community of 11 bacterial strains representing the seven major bacterial phylotypes of the bee gut microbiota [6]. The colonized bees were kept in the laboratory for ten days and fed *ad libitum* either sterile sugar water and pollen (SW+P treatment), or sugar water only (SW treatment). We found a significant difference in gut weight between the two treatments (Fig. 3F, Welch’s T-Test *t*=9.433, P=1.452e-11). While the gut weights of the bees of the SW treatment were comparable to those of forager bees, the gut weights of the bees of the SW+P treatment were markedly higher, exceeding even those of winter bees (Fig. 3D & 3F). We observed a positive correlation between gut weight and microbiota abundance for both the experimentally colonized bees in the laboratory (Fig. 3G) and the conventional worker bees sampled from the hive (Fig. 3E). Moreover, differences in bacterial loads of individual community members between the two experimental treatments mirrored, to a large extent, the differences found between nurses, foragers, and winter bees: most phylotypes were more abundant in bees fed pollen as compared to bees fed sugar water only (Fig. 3H, see **Supplementary Table S4** for statistics). Two exceptions were *Bartonella* and *Frischella.* While *Bartonella* had similar levels between the two experimental treatments (Fig. 3H), its abundance was higher in winter bees and nurses as compared to foragers (Fig. 2A). Notably, *Bartonella* was able to colonize only 75% of all bees when pollen was absent. The dependence on pollen for gut colonization was even more pronounced for *Frischella.* Less than 50% of the experimentally colonized bees of the SW treatment had detectable levels of *Frischella,* and the loads in bees that were colonized were relatively low. In contrast, bees of the SW+P treatment were all colonized and had relatively high and consistent loads of *Frischella* (Fig. 3H).

Taken together, these results show that a pollen diet leads to an increase in gut weight and overall bacterial load providing a plausible explanation for some of the differences in the loads observed between foragers, nurses, and winter bees.

## Discussion

Here, we used a combined approach of qPCR and 16S rRNA gene amplicon sequencing to show that the gut microbiota of adult worker bees markedly differs between nurses, foragers, and winter bees. Nurses and winter bees harbored a larger number of bacteria in the gut than foragers, with most of the dominant community members (except for *Frischella* and *Snodgrassella*) contributing to the increased bacterial loads. Winter bees had the lowest α-diversity of the three bee types, which is explained by the presence of fewer opportunistic colonizers such as *Apibacter, Bombella, or L. kunkeii*. Moreover, a characteristic shift towards high levels of *Bartonella* and *Commensalibacter* was observed in winter bees. These differences in community structure were found across fourteen different colonies and in three different years, suggesting that the “reconfiguration” of the microbiota in winter bees is a conserved feature in colonies in Western Switzerland.

However, regional differences in floral diversity [24] or climate may influence this pattern. Therefore, additional surveys of winter bees in other geographic regions are needed to test for the conservation of this pattern. A recent study carried out in Germany on the effects of winter supplementation feeds found that the relative abundance of certain community members (e.g. *Lactobacillus* Firm-5 and *Bartonella*) increases in winter bees compared to foragers [19], resulting in marked community shifts. The results of this study were also consistent with our findings in that the levels of *Frischella* were significantly lower in winter bees as compared to forager bees. However, this previous study was based on relative abundance data only, i.e. 16S rRNA gene amplicon sequencing.

In contrast to 16S rRNA gene amplicon sequencing, qPCR provides information about the absolute abundance of bacteria and allows determining whether individual community members increase, decrease, or remain the same in terms of bacterial cell number across samples. For example, in our study, the relative abundance of *Snodgrassella* went down in winter bees as compared to foragers and nurses (Fig. 2D). However, this effect was not due to a decrease of the total number of *Snodgrassella*, but rather an increase of other community members as identified by qPCR. In fact, the total abundance of *Snodgrassella* remained the same in foragers, nurses, and winter bees (Fig. 2E). Such quantitative microbiome profiling approaches can reveal important associations between gut bacteria and the host, as previously demonstrated for the human microbiota [32] or the microbiota of caterpillars [50]. We argue that absolute abundances should be routinely assessed when analyzing microbial communities, as changes in absolute abundance – but not necessarily relative abundance – may change the impact of a given bacterium on its environment. Notably, qPCR is a targeted approach, i.e. one can only quantify specific community members for which corresponding primers have been designed, or assess the total amount of bacteria using universal primers. Therefore, a combined approach of qPCR (or any other quantitative method, e.g. flow cytometry) and relative composition analysis (such 16S rRNA gene sequencing or shotgun metagenomics) is preferred, as it provides information about the quantity and directionality of changes in a microbial community.

What drives the observed changes in bacterial loads and community composition in winter bees, nurses, and foragers? A possible explanation could be dietary differences between the analyzed bee types. Foragers mainly feed on nectar and honey, while nurses and winter bees also consume pollen [51]. These dietary differences were also evident in our study, as we found consistent changes in appearance and weight of the dissected guts of foragers, nurses, and winter bees (Fig. 3A-C). Strikingly, our experimental colonization of microbiota-depleted bees with a defined bacterial community showed that pollen in the diet substantially increases gut weight and bacterial loads to levels comparable to those in winter bees. In contrast, bacterial levels in bees fed on sugar water only were more similar to those of foragers (Fig. 3F-H). Therefore, we conclude that diet is an important factor that can explain many of the differences observed between worker bee types. Seasonal changes in gut microbiota composition in wild rodent populations [52] and humans [53, 54] have also been found to coincide with dietary shifts, which is in agreement with the general notion that dietary preferences is the main driver of community differences across a wide range of animals [55–57].

In the case of honey bees, the larger amount of food in the gut is likely to increase the carrying capacity for the gut microbiota. In addition, pollen is a more nutrient-rich diet than nectar, honey, or sucrose offering a larger diversity of different metabolic niches for gut bacteria. Both factors are likely to contribute to the increased bacterial loads in bees fed on pollen as compared to those fed on sugar water only. Recent reports in mice and fly models have shown that an increase in nutritional richness, especially protein quantity, is associated with an increase in overall abundance of the microbiota but a decrease in α-diversity [58, 59]. This is supported by our findings, because we observed an increase in bacterial loads and a decrease in effective number of species in nurses and winter bees that feed on pollen (Fig. 2A and 2B). Consistently, most of the phylotypes that increased in total abundance (*Lactobacillus* Firm5, *Lactobacillus* Firm4, *Bifidobacterium*, *Bartonella*) are located in the rectum, which is the last part of the hindgut where pollen accumulates until bees defecate. In line with this, a previous report showed that the abundances of total bacteria, as well as certain individual phylotypes (*Lactobacillus* Firm-5, *Bifidobacterium*) increase in rectum upon pollen consumption [60]. However, this increase was dependent on the age, and it was not significant when autoclaved pollen was used instead of stored pollen [60]. In contrast to our study, the experimental bees were not inoculated by a defined bacterial community [60], which may greatly impact community growth and dynamics. Overall, despite certain experimental differences, our results seem to be consistent with the data that have been published before.

However, not all changes observed in winter bees could be recapitulated in our colonization experiment. For example, the differences observed *in Bartonella* levels between foragers, nurses, and winter bees (Fig. 2E) were not observed in the experimental bees that were fed with or without pollen (Fig. 3H). Another example is *Frischella*. While this bacterium was less abundant in winter bees than in foragers and nurses, the colonization success of *Frischella* was largely dependent on the presence of pollen in the experimental bees. This suggests that other factors may contribute to community differences found in winter bees as compared to foragers or nurses.

Winter bees have an extended lifespan with an average life expectancy of *∼*6 months as compared to *∼*4 weeks in the case of summer bees (i.e. nurses and foragers)[61]. In contrast, the bees of the colonization experiment were age-matched and sampled ten days after emergence. In the fruit fly, *Drosophila melanogaster*, the physicochemical state of the gut changes with age, resulting in shifts in the composition of microbial communities, mainly characterized by the invasion of certain gut bacterial taxa [62–64]. Therefore, the observed expansion of *Bartonella* and *Commensalibacter* in the gut of winter bees may be related to age. However, despite their old age, winter bees do not display signs of senescence [65, 66] and these differences are likely not due to functional decay in intestinal tissue as reported in flies [64, 67].

Winter bees feed on pollen that has been stored in the hive for several weeks to months. It has previously been shown that the consumption of an aged pollen-diet affects the gut microbiota composition of nurses [12]. It will be important to characterize metabolic differences between the pollen diet of winter and summer bees and to associate such differences with the metabolic capabilities of the different bee microbiota members. For example, *Commensalibacter* and *Bartonella*, the two community members that increased the most in winter bees, carry out aerobic respiration, while most of the other microbiota members are saccharolytic fermenters [22, 68]. Notably, winter bees retain their feces in the gut for extended periods of time, which is likely to affect the physico-chemical conditions and the availability of nutrients in the gut. Moreover, in the absence of defecation, bacteria may accumulate over time in the gut of winter bees, while in nurses or foragers more frequent defecation may result in a faster turnover of the microbiota. Together with differences in the body temperature of bees - in winter it is at ∼21°C and in summer at ∼35°C [69] - this may influence bacterial growth rates. Indeed, in a recent metagenomic study, it was shown that gut bacteria have lower average population replication in old winter bees as compared to young nurse bees, which is indicative of decreased replication rates [18]. Another important point to consider, when carrying our non-culture based community analysis is that these methods usually cannot discriminate between dead and live bacterial cells. Therefore, some of the observed differences could also be attributed to the accumulation of environmental DNA from lysed bacterial cells.

Finally, winter bees show reduced expression of immune genes [70–73], and have an altered protein metabolism [74] as compared to nurses and foragers, another factor which may influence the total bacterial loads and the community composition in the gut.

In the case of *Frischella,* it is tempting to speculate that the decrease in colonization levels in old winter bees may be a consequence of the specific immune response elicited by the host towards this bacterium [10], eliminating it from the gut as the bee ages. In the case of *Snodgrassella*, it is interesting to note that the levels of *Snodgrassella* barely changed across worker bee type or the two diet treatments in the experiment. This suggests that the colonization of *Snodgrassella* is not modulated by the dietary state, the physicochemical conditions in the gut, or the abundance of other community members. A possible explanation could be that the niche of *Snodgrassella* is dependent on the host rather than the diet, because it selectively colonizes the epithelial lining of the ileum, which presents a physically restricted niche [8, 75, 76].

Beside the increase of *Commensalibacter* and *Bartonella*, another intriguing characteristic of the winter bee gut microbiota was the disappearance of minor, non-core community members in the bee gut microbiota. We can exclude that these differences in community composition are due to a community sampling bias, because nurses had similar bacterial loads as winter bees, but showed the opposite trend in respect to the presence of minor community members. We hypothesize that these minor community members are transient colonizers that cannot persist in the bee gut environment over longer periods of time and hence disappear in old winter bees. As some of these bacteria, e.g. *Serratia or Klebsiella,* present potential pathogens of bees, there may be also mechanisms in place that increase colonization resistance against such opportunistic colonizers in winter bees. Moreover, during the foraging season adult worker bees are more likely to pick up environmental bacteria from e.g. flowers, facilitating their dissemination in the hive environment during the summer but not in winter.

Most of the recent colony losses have occurred during the winter months [33, 34]. Consequently, winter bees are highly critical for colony survival and a better understanding of the factors influencing their health status - including the gut microbiota - is needed. In summary, our analysis revealed that the gut microbiota of winter bees undergoes characteristic shifts. These changes may have important consequences for the host. Therefore, future studies should specifically focus on the functional role of the gut microbiota in winter bees, and colony health.

## Acknowledgements

We would like to thank Paul Reymond, Clément Etter, Katherine Lane for their help during the sampling of honey bees. We are also grateful to Andrew Quinn and Fabienne Wichmann for comments on the manuscript. This study was supported by the European Research Council (ERC-StG ‘MicroBeeOme’) and the Swiss National Science Foundation (grant number 31003A_160345 and 31003A_179487) received by PE, and a Marie Skłodowska-Curie Postdoctoral Fellowship (grant agreement ID 797113) to JL. The funder had no role in study design, data collection and analysis, decision to publish, or preparation of the manuscript.

## Data accessibility

All scripts and datasets will be deposited to Zenodo upon acceptance. For the revision they can be found on: https://drive.switch.ch/index.php/s/zHsqyMOztvAr8Vu.

Sequencing data is deposited on NCBI under BioProject ID PRJNA578869.

## Supplementary Material

**Supplementary Figures:** Contains Figures S1-S3

**Supplementary Table S1:** Primers used in this study and stndard curve characteristics.

**Supplementary Table S2:** Details of statistics used in Fig 1.

**Supplementary Table S3:** Details of statistics used in Fig 2.

**Supplementary Table S4:** Details of statistics used in Fig 3.

**Supplementary Table S5:** Details of statistics used in Supplementary Fig S5.

**Supplementary Table S6:** Data related to the number of reads obtained from 16S rRNA gene sequencing and after each step of filtration.

**Supplementary Table S7:** Number of rRNA gene loci for each taxonomic group

