## Supplementary Figures for "Gut microbiota structure differs between honey bees in winter and summer"

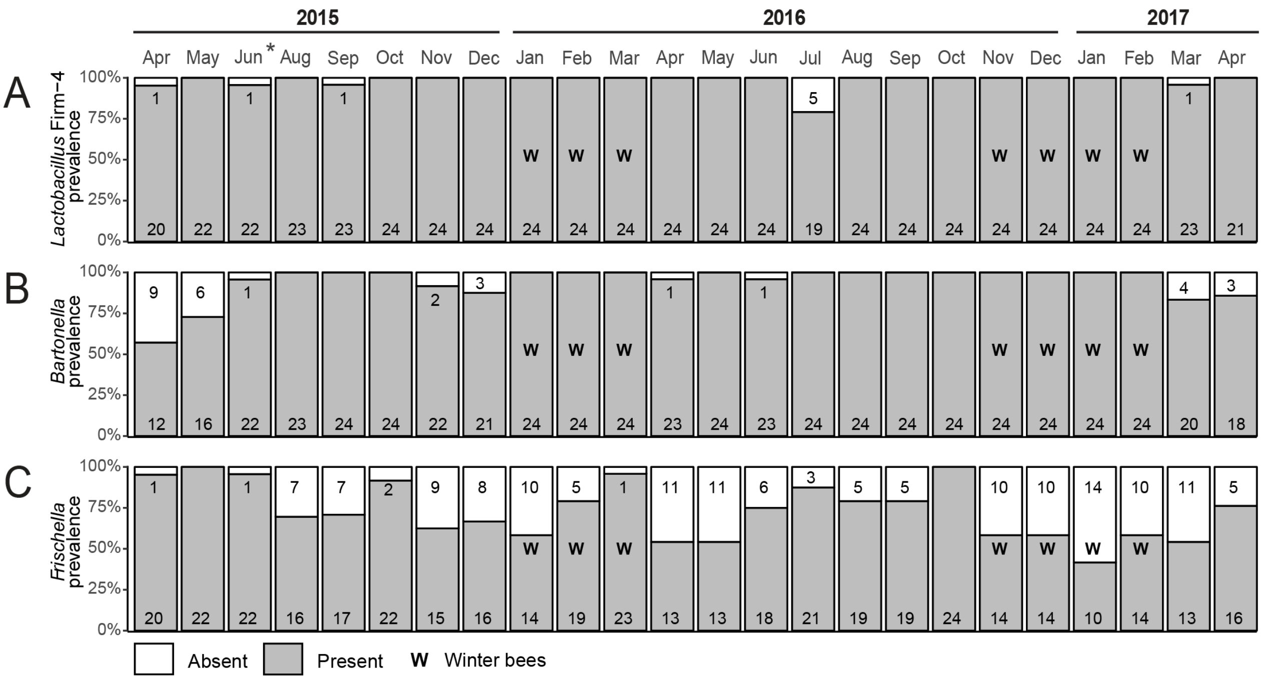


**Supplementary Fig. S1. Prevalence of phylotypes that were not detected in every honey bee of the hive sampled over two years.** Monthly prevalence (i.e. the proportion of bees in which the bacterial phylotype was detected in a given month) of **(A)** *Lactobacillus* Firm-4, **(B)** *Bartonella* and **(C)** *Frischella*. The targeted bacterium was considered absent when the qPCR signal was below the detection limit of the primer set (see **Supplementary Table S1**). The numbers in the upper and lower part of the bar plots indicate the number of bees in which the target is absent and present, respectively. Asterisk indicates missing data for July 2015 due to a sampling/extraction error.

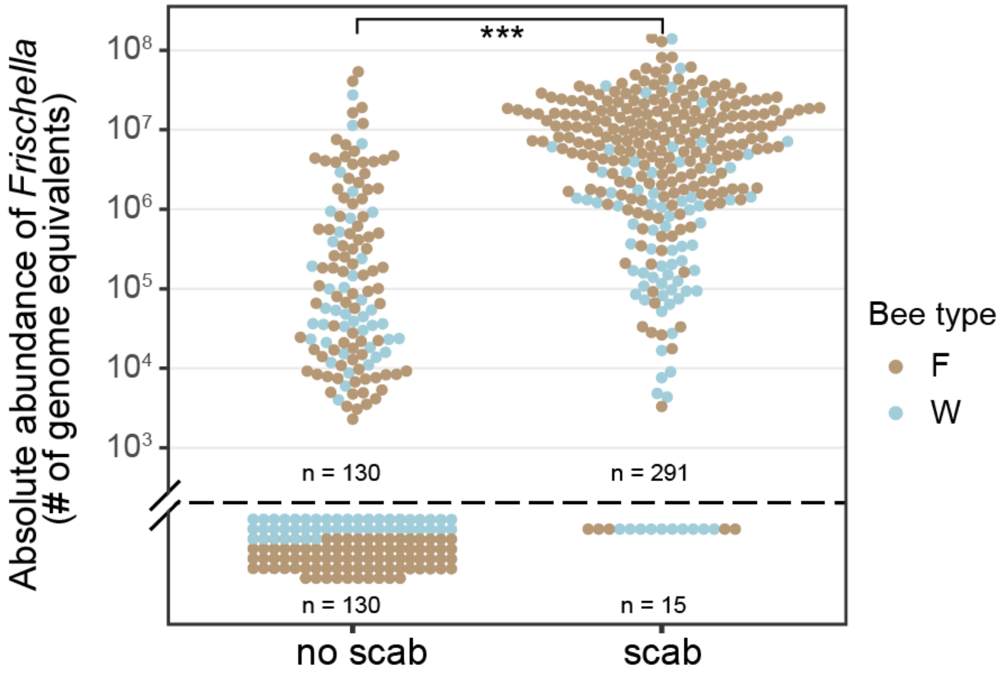


**Supplementary Fig. S2. *Frischella* loads correlate with scab presence.** *Frischella* normalized number of cells (see methods) in guts of foragers (F) and winter bees (W) with or without scab phenotype in the pylorus region. Data points under the black dashed line correspond to samples with no detectable level of *Frischella* as based on the detection limit of our qPCR method. Statistics: permutation T-Test. ***, P < 0.001.

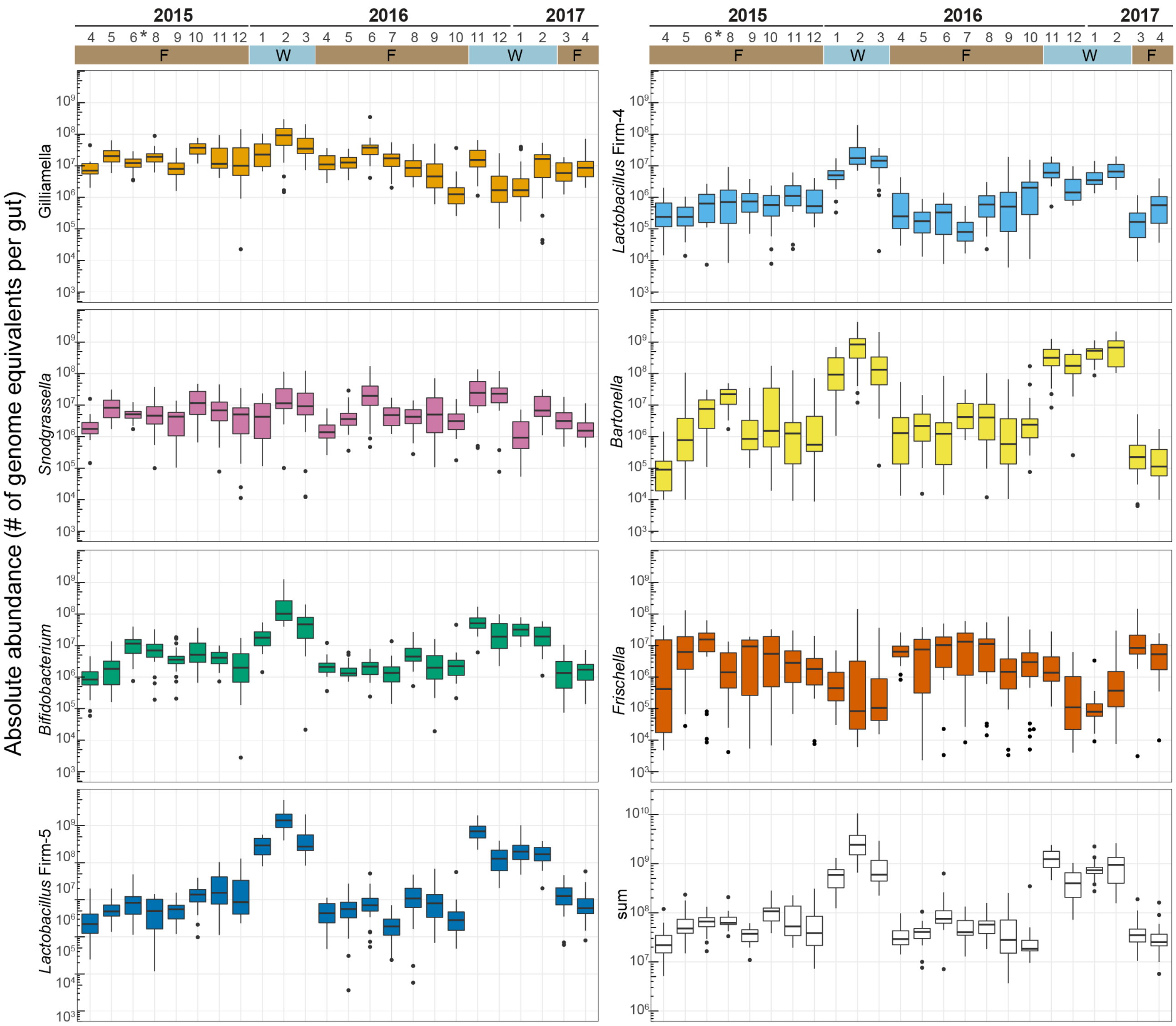


**Supplementary Fig. S3. Monthly abundances of seven core microbiota members monitored in a single hive over two years.** Absolute abundance of the seven monitored phylotypes in individual foragers and winter bees sampled over two years: *Gilliamella*, *Snodgrassella*, *Bifidobacterium*, *Lactobacillus* Firm-5, *Lactobacillus* Firm-4, *Bartonella*, and *Frischella*. The eighth plot shows the summed abundances of each bee. Only bees with detectable levels were considered. Asterisk indicates missing data for July 2015 due to a DNA extraction error.

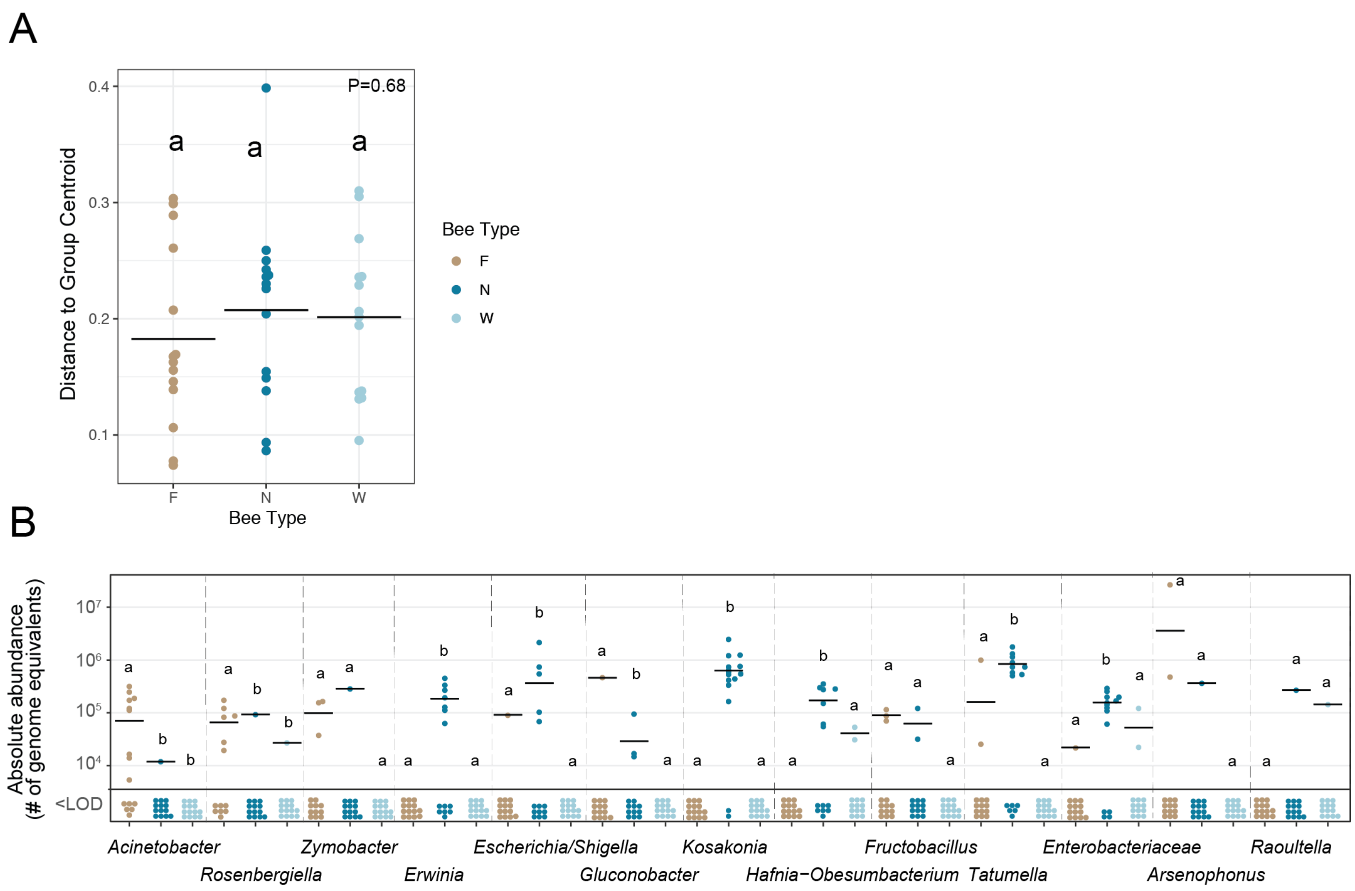


**Supplementary Fig. S4. (A)** Dispersal of communities in each bee type based on Bray-Curtis distances. Symbols represent Euclidean distances of communities to group centroid. Same letters indicate lack of significance based on Beta-dispersal Permutest (See **Supplementary Table S3** for statistics) (**B**) Absolute abundance of the remaining community members (i.e. not presented in **Fig. 3F**) in foragers (F), nurses (N), and winter bees (W) across hives, as determined based on the number of genome equivalents per gut calculated by multiplying the relative abundance of each phylotype by the total 16S rRNA gene copy number. <LOD, below limit of detection of the 16S rRNA amplicon sequencing, i.e. no reads were obtained for that particular taxa in the respective sample. Different letters indicate statistically different group based Permutation ANOVA (B) followed by Tukey HSD Test. For detailed statistics **see**

**Supplementary Table S3.** The black bar represents the mean.

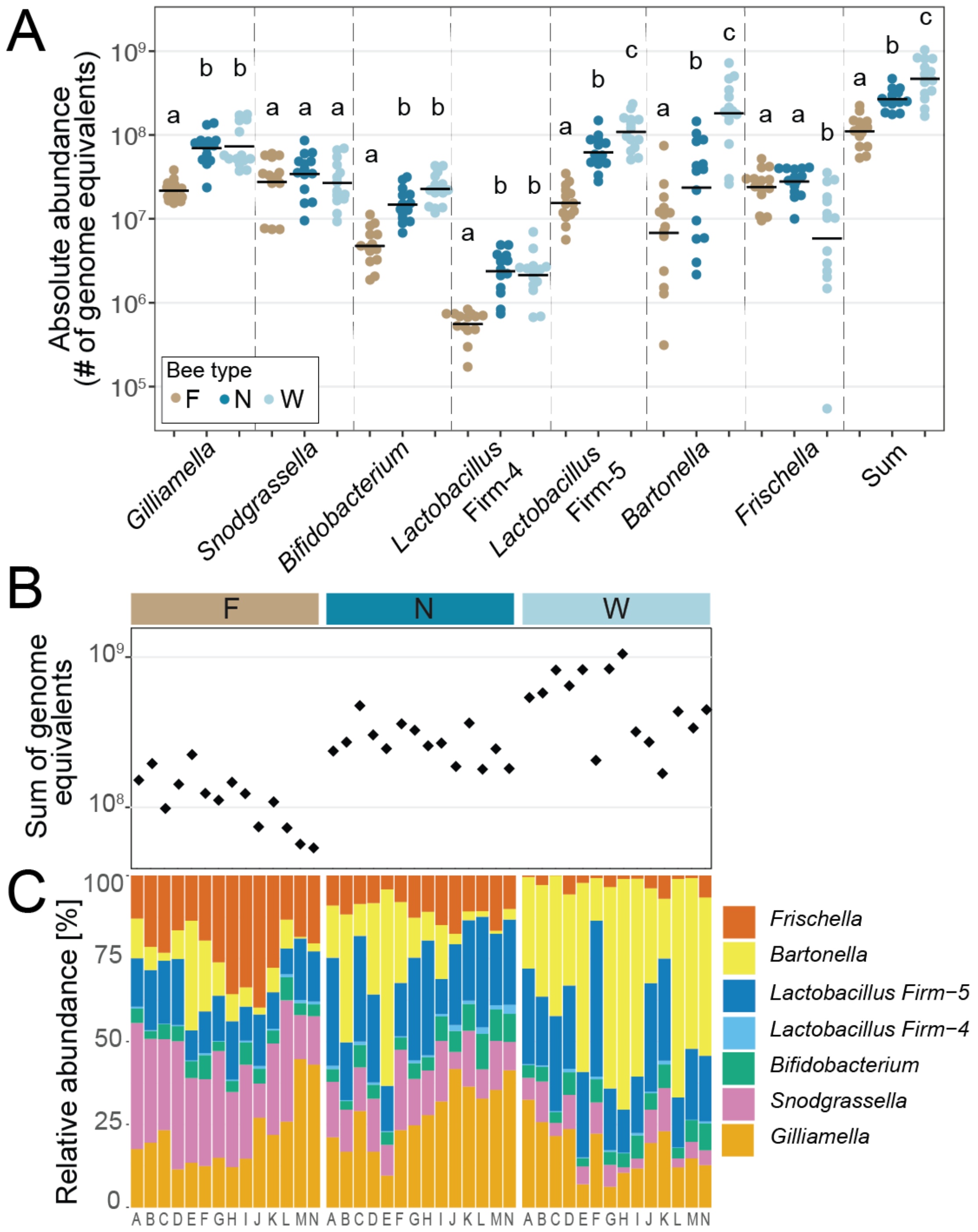


**Supplementary Fig. S5. Bacterial loads and community composition between foragers, nurses, and winter bees across 14 colonies determined by qPCR using phylotype-specific primers. (A)** Absolute abundances, i.e. genome equivalents per gut, of the seven dominant phylotypes in foragers (F), nurses (N), and winter bees (W)

across hives, as determined by qPCR with phylotype-specific primers. The sum of the abundances of the seven monitored phylotypes is also given. Mean values are shown as black horizontal lines. (**B**) Black diamonds indicate the sum of the total abundances of the seven monitored phylotypes in each sample corresponding to the samples shown in the panel below. (**C**) Relative community composition of the gut microbiota based on the abundances of the seven monitored phylotypes. Capital letters below the stacked bars indicate the hive of origin of each sample. In panel A, levels (bee types) not connected by the same letter are significantly different (Permutation ANOVA followed by Tukey HSD test, See **Supplementary Table S5** for detailed statistics).
